# The flexibility of ACE2 in the context of SARS-CoV-2 infection

**DOI:** 10.1101/2020.09.16.300459

**Authors:** E. P. Barros, L. Casalino, Z. Gaieb, A. C. Dommer, Y. Wang, L. Fallon, L. Raguette, K. Belfon, C. Simmerling, R. E. Amaro

## Abstract

The COVID-19 pandemic has swept over the world in the past months, causing significant loss of life and consequences to human health. Although numerous drug and vaccine developments efforts are underway, many questions remain outstanding on the mechanism of SARS-CoV-2 viral association to angiotensin-converting enzyme 2 (ACE2), its main host receptor, and entry in the cell. Structural and biophysical studies indicate some degree of flexibility in the viral extracellular Spike glycoprotein and at the receptor binding domain-receptor interface, suggesting a role in infection. Here, we perform all-atom molecular dynamics simulations of the glycosylated, full-length membrane-bound ACE2 receptor, in both an apo and spike receptor binding domain (RBD) bound state, in order to probe the intrinsic dynamics of the ACE2 receptor in the context of the cell surface. A large degree of fluctuation in the full length structure is observed, indicating hinge bending motions at the linker region connecting the head to the transmembrane helix, while still not disrupting the ACE2 homodimer or ACE2-RBD interfaces. This flexibility translates into an ensemble of ACE2 homodimer conformations that could sterically accommodate binding of the spike trimer to more than one ACE2 homodimer, and suggests a mechanical contribution of the host receptor towards the large spike conformational changes required for cell fusion. This work presents further structural and functional insights into the role of ACE2 in viral infection that can be exploited for the rational design of effective SARS-CoV-2 therapeutics.

**Statement of Significance:** As the host receptor of SARS-CoV-2, ACE2 has been the subject of extensive structural and antibody design efforts in aims to curtail COVID-19 spread. Here, we perform molecular dynamics simulations of the homodimer ACE2 full-length structure to study the dynamics of this protein in the context of the cellular membrane. The simulations evidence exceptional plasticity in the protein structure due to flexible hinge motions in the head-transmembrane domain linker region and helix mobility in the membrane, resulting in a varied ensemble of conformations distinct from the experimental structures. Our findings suggest a dynamical contribution of ACE2 to the spike glycoprotein shedding required for infection, and contribute to the question of stoichiometry of the Spike-ACE2 complex.

## Introduction

Angiotensin-converting enzyme 2 (ACE2) acts as the extracellular receptor for the severe acute respiratory syndrome coronavirus 2 (SARS-CoV-2) (1–3), the virus responsible for the COVID-19 pandemic that has catastrophically affected the world since its first identification in December 2019 (4–7). ACE2 is a membrane protein found in lungs, kidneys, heart and intestine cells (8, 9) that plays a physiological role in cardiovascular regulation via the cleaving of intermediates in the maturation process of angiotensin, a peptide hormone involved in vasoconstriction control (10–14). ACE2 is a homodimer with a large claw-like extracellular head domain, a small transmembrane domain and a short intracellular segment (8). The head can be further subdivided into the catalytic zinc-binding peptidase domain (PD, residues 19 to 615) (15), and the smaller neck domain (residues 616 to 726), which is where the majority of the homodimer interactions seems to lie (16). The neck domain is further connected to the single-helix transmembrane (TM) domain by a long linker (Figure 1a). ACE2 can also function as a membrane-trafficking chaperone for B^0^AT1, an amino acid transporter (17), and it was in fact only in complex with this partner that the single TM helix of ACE2 could be resolved (16).

**Figure 1.**
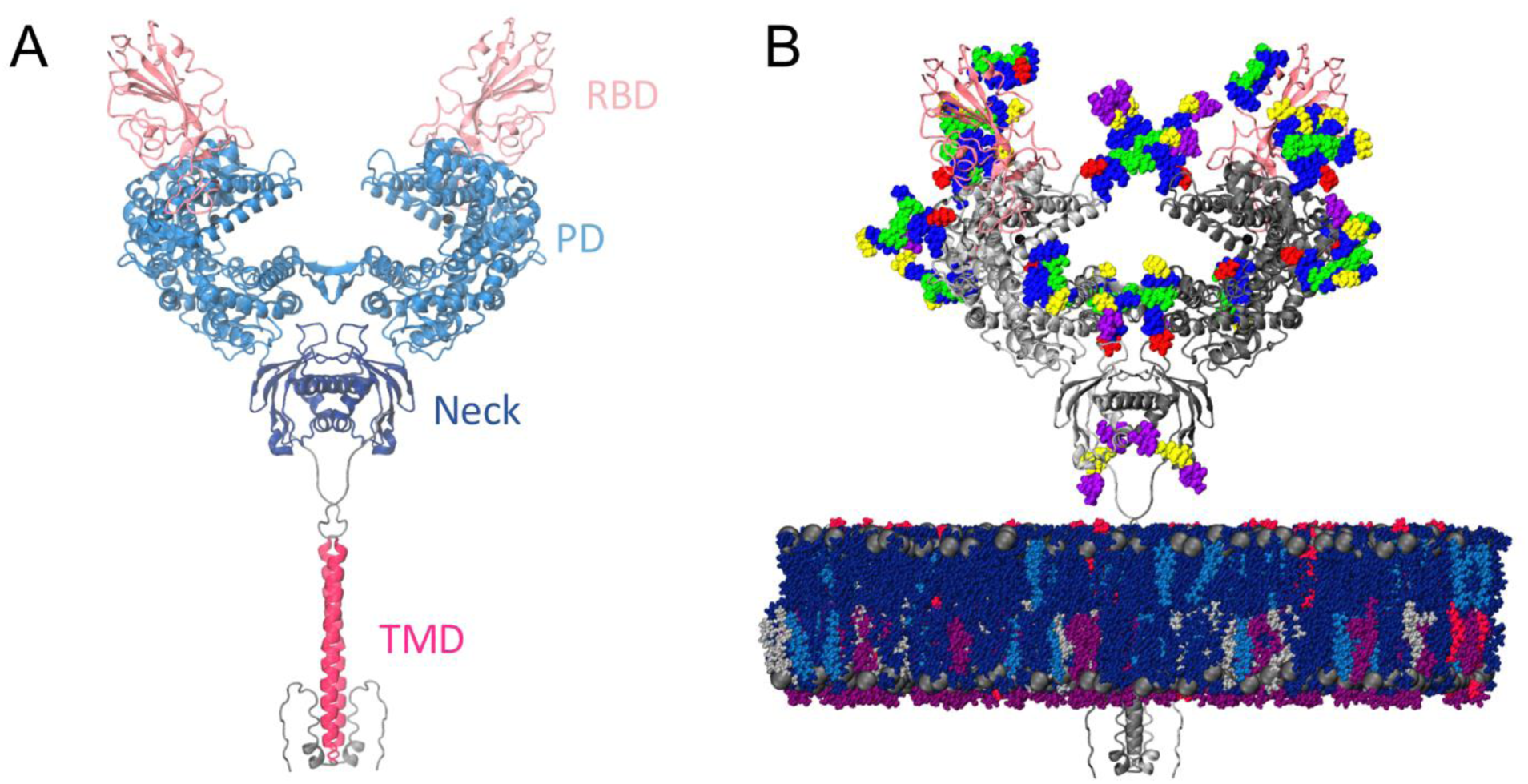
Model structure. **(a)** Full-length ACE2 homodimer protein structure in complex with spike protein RBDs. ACE2 peptidase, neck and transmembrane domains are shown with cartoons highlighted in blue, navy and magenta, respectively. Spike RBDs are depicted with pink cartoons. **(b)** Fully glycosylated and membrane-embedded model. ACE2 and RBDs are represented with gray and pink cartoons, respectively. Atoms of N-/O-glycans are shown with per-monosaccharide colored spheres, where GlcNAc is highlighted in blue, mannose in green, fucose in red, galactose in yellow, and sialic acid in purple. Lipid heads (P atoms) are represented with grey spheres, whereas lipid tails are depicted with a licorice representation using the following color scheme: POPC (navy), POPI (violet), POPE (silver), CHL (blue), PSM (magenta).

SARS-CoV (18) (responsible for 8096 cases in 2002 (19)) and now the closely related SARS-CoV-2 hijack ACE2 as the host cell receptor to its large extracellular spike (S) glycoprotein (1, 20). The spike’s receptor binding domain (RBD) in the “up” conformation binds to ACE2’s PD with high affinity (21), and the resolved ACE2-RBD complex consists of a dimer of heterodimers, with each monomer of the ACE2 homodimer interacting with one RBD (thus forming a heterodimer, Figure 1a). Cellular recognition and binding to ACE2’s peptidase domain via the RBD is proposed to initiate a series of complex conformational transitions in the S homotrimeric protein, leading to the shedding of its S1 subunit and fusion to the host cell membrane driven by the S2 subunit (22–26), ultimately resulting in the infection of the host cell. Downregulation of ACE2 and accumulation of angiotensin II due to spike binding is also associated with acute respiratory distress syndrome (ARDS) and acute lung failure (27–31), contributing to SARS-associated symptoms. As such, the S glycoprotein and ACE2-S complex are considered key targets for drug and antibody development efforts aiming to curtail the virus’ remarkable transmissibility and negative effect on human health (1, 32–35), including exploiting the ACE2-S high affinity with recombinant soluble ACE2-antibody constructs (35–38).

Experimental and biophysical studies of the SARS-CoV-2 RBD and soluble ACE2’s PD complex have suggested structural factors likely responsible for the higher affinity and infectivity of SARS-CoV-2 compared to SARS-CoV (39–41), and revealed significant dynamics at the RBD-PD interface (41, 42). Additionally, cryoEM and computational studies of the full-length spike glycoprotein have recently suggested a significant degree of flexibility of the spike’s stalk and at the ACE2-RBD interface (43, 44), evidencing the need to study these macromolecular complexes in the context of the cell surface instead of in static, cryogenic conditions. Here, we perform all-atom molecular dynamics (MD) simulations of the full length, membrane-embedded and glycosylated ACE2 homodimer both in the apo state and in complex with RBD to study the molecular origins of the ACE2-S flexibility on the host receptor side. Seven complex N-glycans and one O-glycan in ACE2 were modeled according to glycoanalytic data (45–47), as well as glycan N343 in RBD for the RBD-bound simulations (Figure 1b). The B^0^AT1 transporter solved in the cryo-EM structure was not included in our simulations in order to inform on the intrinsic dynamics of ACE2. B^0^AT1 is mainly expressed in kidneys and intestines (48), whereas ACE2 can also be found in lungs and heart tissues, supporting the likelihood that ACE2 can be found un-complexed with B^0^AT1 upon cellular recognition and binding to S.

The simulations reveal an exceptional structural plasticity of the full-length ACE2 homodimer, pinpointing a large tilting of the head relative to the TM domain, as well as profuse mobility of the TM helix in the membrane. Remarkably, the homodimer interface at the level of the neck domains remains stable despite the dramatic motions, as well as the ACE2-RBD contacts, emphasizing the high affinity interaction between them. A systematic characterization of glycan-protein and glycan-glycan contacts indicates a possible role of glycan N53 in both homodimer and heterodimer interactions. Overall, the RBD does not seem to significantly affect the dynamics of ACE2 compared to the apo state, although that might differ in the presence of the full-length spike. Taken together, the remarkable ACE2 flexibility indicates a mechanical contribution to the S1/S2 conformational changes required for cellular fusion and infection, and suggests the structural basis for the possibility of finding two or more ACE2 complexes bound to the same S glycoprotein with two or more “RBD-up” conformations.

## Methods

### ACE2 system construction

Coordinates of the ACE2-RBD complex were taken from the full length cryo-EM structure, PDB ID 6M17 (16), removing the coordinates from the co-complexed B^0^AT1 dimer. Missing C terminal residues of the ACE2 transmembrane helices were modeled using I-TASSER (49–51) based on the known sequence (residues 769 to 805), while missing N terminal residue coordinates (residues 19 to 21) were copied from 6M0J following alignment of the N terminal helix. Zinc coordinating residues and a coordinating water molecule were taken from 1R42 as the zinc coordination site is poorly resolved in 6M17.

ACE2 and RBD glycosylation was defined according to glycoanalytic data (45, 47) and modeled using the *Glycan Reader & Modeler tool* (52) integrated into *Glycan Reader* (53) in CHARMM-GUI (54). In total, 7 complex, bi-antennary N glycans and 1 O glycan were added to ACE2, as well as 1 N glycan to RBD (Supplementary Table 1). Only one O glycan was included at site 730 as analytic data suggest extremely low stoichiometry at the other O glycosylation sites (46). The apo ACE2 model was created by deleting the RBDs of the complete ACE2-RBD model.

### Membrane modeling

The plasma membrane modeled in this study was composed of 56% POPC, 20% CHL, 11% POPI, 9% POPE, and 4% PSM. The lipid composition was estimated based on the known lipid compositions of mammalian cellular membranes (55, 56). It is hypothesized that phospholipids containing charged headgroups such as PI and PS are more likely to face the cytoplasmic side of the membrane and are additionally thought to aid in tolerance of increased membrane curvature (55). Using a precedent set by a 2014 coarse-grained molecular dynamics study of the asymmetrical mammalian plasma membrane, the lipids were partitioned according to the outer *versus* inner leaflet enrichment factors of 2.0, 1.2, 0.0, 0.25, and 2.0 for POPC, CHL, POPI, POPE, and PSM, respectively (56). To reduce the chemical complexity of the system for simulation purposes, PS lipids were not included in these calculations. The small percentage (4%) of PS recorded in the literature is represented in the membrane by PI lipids.

An asymmetric 350 Å x 350 Å lipid bilayer according to the above specifications was generated using CHARMM-GUI’s input generator (54). The lipids were packed to an approximate equilibrium area per lipid of 63 Å^2^. Prior to insertion of ACE2 and subsequent trimming, the membrane patch contained a total of 2,432 POPC, 870 CHL, 460 POPI, 404 POPE, and 128 PSM lipids.

### System preparation and molecular dynamics simulations

Histidine protonation states at pH 7.0 were verified using PROPKA on Maestro (Schrödinger, LLC, New York, NY). The models were parametrized using PSFGEN and CHARMM36 all-atom additive force fields for protein, lipids, and glycans (57), fully solvated in TIP3P water boxes (58) with 150 mM NaCl. The total number of atoms is 738,696 for the apo system (size: 18.7 nm × 18.9 nm × 23.7 nm) and 783,954 for the RBD-bound system (size: 18.7 nm × 18.9 nm × 25.1 nm).

Minimization, equilibration and production simulations were performed on the Frontera computing system at the Texas Advanced Computing Center (TACC) using NAMD 2.14 (59), as described in detail in Casalino *et al* (60). Each system was run in triplicates for 1μs each.

### ACE2 angles and distances calculations

To quantify the range of motion of ACE2 in the simulations, several angle and distance metrics were developed. Calculation was performed using MDTraj (61) with visualization through VMD (62). The 6M17 cryo-EM structure was used as the reference structure.

Head tilt angle relative to the transmembrane domain was calculated by first aligning the dimer’s coordinates to the reference cryo-EM TM domains, and angle calculated between the centers of mass of the reference’s dimer peptidase domains (residues 18 to 600), reference’s TM helices (residues 747 to 774), and monomer’s PD at each frame in the simulation. Helix tilt angle was computed as the angle between a vector defining the membrane’s normal and a vector connecting residues 741 and 765 at the extremities of the helix.

Revolution angle was calculated between the center of mass of the monomer’s PD in the reference conformation, the center of mass of the reference’s TM domain, and the center of mass of the monomer’s PD at each frame in the simulation following alignment of the monomer’s TM helices. Buckling angle was calculated using the *xy* projections of the center of mass of monomer’s A PD at frame *f*, the center of mass of the reference dimer’s PDs, and center of mass of monomer’s B PD at frame f, after alignment of the trajectories to the reference dimer neck domains (residues 617 to 726).

Distance between the monomer’s head domains in the homodimer was calculated by determining the distance between each monomer’s PD center of mass. Distance between the head domain and membrane corresponds to the minimum distance between the PD’s heavy atoms and membrane’s phosphorous atoms at each frame of the simulation. Distance between TM helices was calculated based on the distance between their centers of mass.

### Fraction of native contacts and glycan contacts

Fraction of native contacts was calculated according to Mehdipour and Hummer (63). The 6M17 cryo-EM structure was used as the reference structure for identification of native contacts. The ACE2-RBD interface was subdivided into three interacting regions according to the interacting residues pairs listed on Supplementary Table 2.

A systematic characterization of contacts established by each glycan in the system was performed using MDTraj (61), using a cutoff of 3.5 Å between the heavy atoms.

### S model construction

The spike model was obtained from our previous simulations (60). The simulations included only the solvated spike. All atoms except for the spike protein and glycans were removed, along with the lower part of the stalk region of each protomer (residues 1165-1273). Since the cryo-EM model was missing density for portions of the RBD, we replaced the RBD coordinates (residues 355 to 494 for closed RBD, 339 to 523 for open RBD) of each protomer in both models with the RBD coordinates from the crystal structure of the RBD bound to ACE2 (PDB 6M0J (64)). The RBD structure from 6M0J was aligned with the backbone heavy atoms (alpha-carbon, carbonyl-carbon, and nitrogen) of each RBD in the initial spike model. We then grafted the RBD coordinates onto the spike at the hinge region, which resolved missing loops as well as introducing a disulfide bond in the RBD. The remaining disulfides not resolved in the cryoEM structures were assessed based on distance criteria and sequence conservation. The system was built using the *ff14SBonlysc* (65) and GLYCAM (66) force fields for the protein and glycan atoms, respectively. These were explicitly solvated in OPC3 water (67) with a 200 mM NaCl buffer (68). The RBD-up and -down systems both consisted of 1,298,646 atoms, and were simulated on Frontera at TACC, and SDCC at BNL using the *pmemd*.*CUDA* module of Amber20. The spike systems were equilibrated using a 10-step protocol. First, the water molecules were minimized for 1,000 steps using steepest descent, and then for an additional 9,000 steps with conjugate gradient while the rest of the system was positionally restrained with 1 kcal/(mol * Å^2^) restraints. The systems were then heated to 310 K at constant volume, again with all atoms except hydrogens and waters restrained with 100 kcal/(mol * Å^2^) positional restraints. The box size and density were then equilibrated over 1 ns with constant pressure, with the same positional restraints as the previous step. The restraints were then lowered to 10 kcal/(mol*A) for an additional 1 ns of equilibration, before a second minimization. This minimization consisted of 10,000 steps of conjugate gradient with positional restraints now applied only to only backbone atoms (alpha-carbon, carbonyl-carbon, and nitrogen), using a force constant of 10 kcal/(mol * Å^2^). The next three steps of equilibration were MD for 1 ns each at constant NPT with positional restraints on protein backbone atoms at 10, 1, and 0.1 kcal/(mol * Å^2^), respectively. This was followed by a final 1 ns of unrestrained MD at constant NPT before beginning production.

To generate a 3-up model of the prefusion spike protein, steered molecular dynamics (SMD) was used as implemented in the Amber NFE toolkit (69). The initial structure was an all-closed model from the equilibration described above. To generate a structure used as reference for steering, the open monomer from the 1-up model was aligned to the other two closed monomers by overlapping S2 domains. The closed RBD domains were then replaced with the open model of the aligned monomer. No equilibration was performed on the resulting reference structure since it was not subjected to simulations, and only used as a steering target. Root mean squared deviation (RMSD) was used as the collective variable (CV) to guide SMD. A separate CV was used for the opening of each RBD. In each CV, the RMSD region includes the RBD (residue 338 - 517) and 3 helices in S2 (residue 747 – 782, 946 – 966, 987 - 1034). The 3 RMSDs were gradually decreased by SMD from their initial values to 0 during 20 ns of simulation time at 310K in the NVT ensemble, with a 4fs time step enabled by hydrogen mass repartitioning (70), using a spring constant of 10000 kcal/mol/Å^2^. Weak (1 kcal/mol/Å^2^) positional restraints were applied to the S2 helices, which were relatively stable during SMD simulations.

## Results

### The ACE2 dimer shows pronounced flexibility

Simulations of RBD-bound and apo ACE2 evidenced a striking degree of flexibility in the ACE2 homodimer. With respect to the fairly vertical, extended conformation of the initial cryo-EM structure (16), the most striking fluctuation observed during the simulations is characterized by a tilt of the head relative to the long axis of the respective monomer’s transmembrane helix. While each monomer in the reference cryo-EM structure displays a tilt angle of 16°, structures in the simulations sample tilt angles that range from 0° to 50° (Figure 2a and d). This tilt motion, combined with an overall “shrinking” of the initial extended conformation, moves the head towards the membrane, with head-membrane distances varying from 30 to 84 Å and the great majority of conformations (98% and 98.6% of the frames of apo and RBD-bound simulations, respectively) exhibiting the head domain closer to the membrane than the starting cryo-EM structure (Figure 2b). The presence of the RBD does not seem to affect the dynamics, with apo and RBD-bound simulations showing average head-membrane distances of 59.1±6.9 Å and 56.5±8.8 Å, respectively.

**Figure 2.**
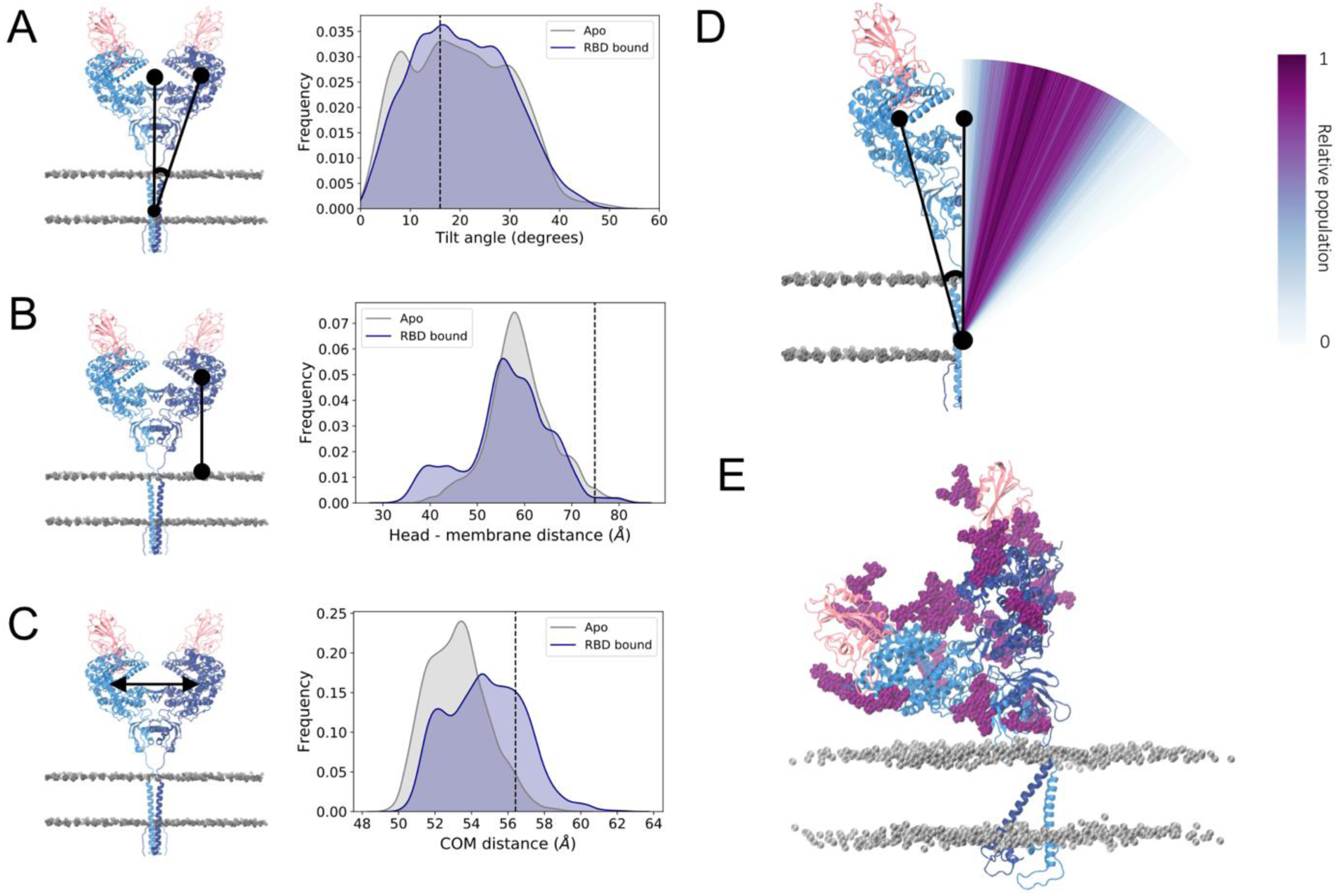
Tilt motion of ACE2. **(a)** Head tilt angle distribution relative to the transmembrane domain long axis for apo (grey) and RBD-bound (navy) simulations. The angle value corresponding to the cryo-EM conformation is indicated by a black line. Left panel shows a representation of the metric, with ACE2 monomers colored dark and light blue, RBDs colored pink and phosphorus atoms from membrane’s lipid heads shown in grey in van der Waals representation. **(b)** Distribution of minimum distance between PD’s center of mass and membrane. **(c)** Distribution of the ACE2 monomer heads’ center of mass distance. **(d)** Visual representation of the tilt angle distribution for the RBD-bound simulations with a color gradient according to the relative population. **(e)** Example of a highly-tilted ACE2 homodimer conformation sampled in the simulation. ACE2 and RBD glycans shown in dark purple.

Remarkably, the head tilt motion occurs in a concerted fashion between the monomers, such that as one monomer bends towards the membrane, with large tilt angle values, the other monomer follows this deformation by adopting a more extended conformation with lower angle values (Figure 2e, Supplementary Figure 1). Accordingly, the distance between the head domains fluctuates only slightly, varying by no more than 6 Å (Figure 2c) and resulting in a stable relative position of the heads within the homodimer. The majority of the conformations display the two heads slightly closer to each other than in the resolved cryo-EM structure, while the presence of the spike RBD shifts the distribution slightly towards more open conformations.

In concert with the tilt motion described above, the ACE2 head also undergoes displacement in the *xy* plane around the long axis of the TM helix, as shown for one of the replicas in Figure 3a. Computation of each monomer’s revolution angle suggests a twisting of the flexible linker that connects the neck to the TM domain, with several significant alternative conformations exhibiting almost 180 degrees rotation of the head from its starting position (Figure 3b). It is important to highlight that the head revolution is measured here for each monomer independently, following an alignment of that monomer’s transmembrane domain. As the monomer’s TM helices are not in contact with each other and thus can move independently in the membrane (Figure 4a), the twisting motion of one of the monomers is not necessarily accompanied by an equivalent twist of the other monomer, avoiding a twist of the flexible linkers around each other. Instead, visual observation indicates that the other monomer revolves as a whole around the transmembrane helix of the twisting monomer (Supplementary Figure 2), keeping the head dimer interface overall intact. Thus, despite this pronounced motion, the other monomer follows the twist by retaining the heads’ symmetry, and the relative angle between the heads in the *xy* plane remains close to the initial 180° (Figure 3c).

**Figure 3.**
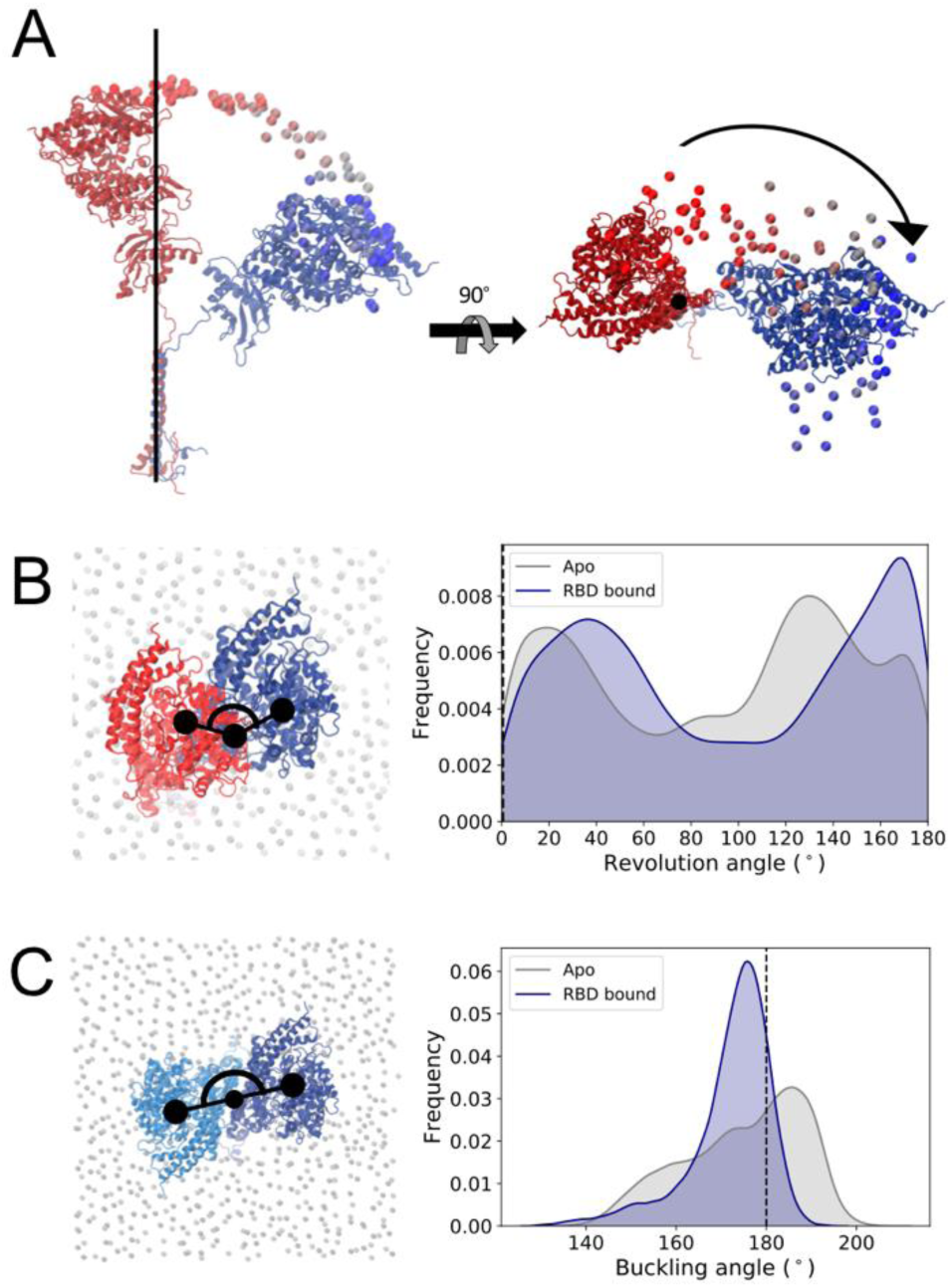
ACE2 revolution relative to a plane perpendicular to the transmembrane helix’s long axis. **(a)** Representation of a monomer’s degree of flexibility in one of the replicas, showing the time evolution of the position of the Cα atom of Gln325 colored from dark red (t=0) to dark blue (t=1000 ns). Conformations aligned to the cryo-EM’s reference TM domain Cα atoms shown in van der Waals representation, initial and final monomer conformations shown in cartoon representation. **(b)** Head revolution angle distribution for apo (grey) and RBD-bound (navy) simulations. The angle value corresponding to the cryo-EM conformation is indicated by a black line. Left panel shows representation of the metric, with monomer’s head initial position shown in red, the same monomer at a time t in dark blue, and phosphorus atoms from membrane’s lipid heads shown in grey in van der Waals representation. **(c)** Relative orientation of the monomer’s head in the heterodimer. ACE2 monomers colored dark and light blue.

**Figure 4.**
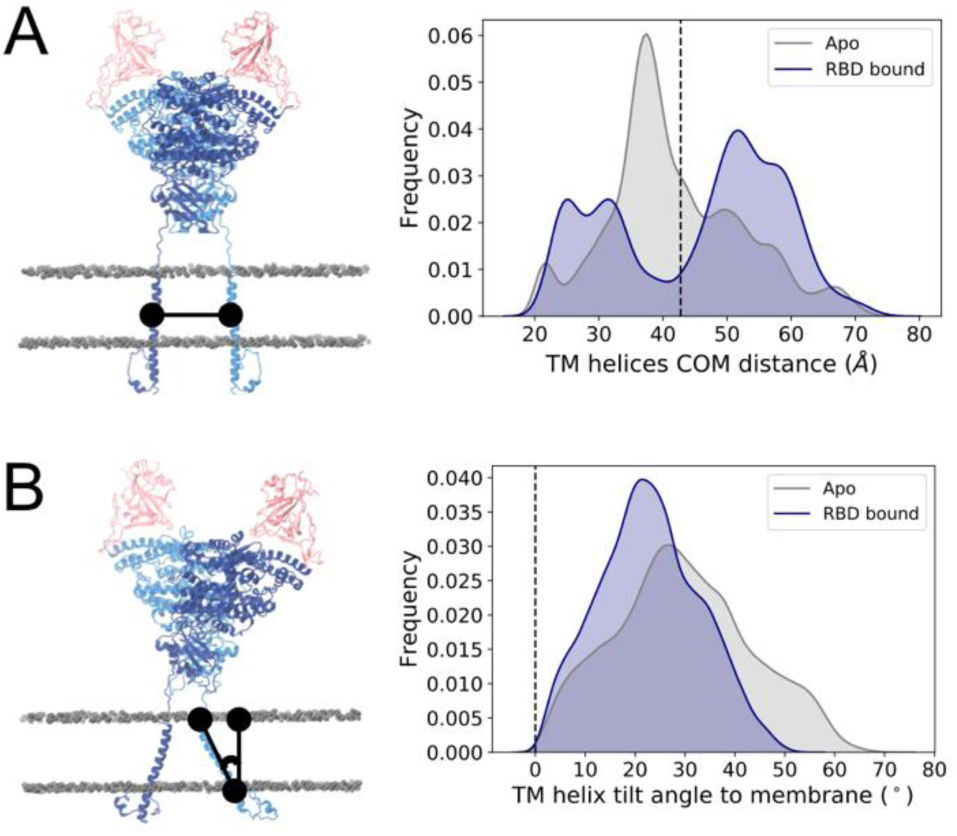
Transmembrane helix dynamics. **(a)** Distance between the center of mass of each monomer’s TM helix. **(b)** TM helix tilt angle relative to the membrane’s normal. Values corresponding to cryo-EM conformation are indicated by a black line. Left panels show representation of the metric, with ACE2 monomers colored dark and light blue, RBDs colored pink and phosphorus atoms from membrane’s lipid heads shown in grey in van der Waals representation.

The simulations indicate that the conformational variability of ACE2 occurs not only due to flexibility at the linker connecting the transmembrane and head domains, but also due to motions of the transmembrane helix in the membrane. In contrast to other multimeric transmembrane domains such as the coiled coil trimer of S (21, 26, 60), each ACE2 monomer is anchored to the membrane by a single helix, which does not interact with that of the opposing monomer (Figure 4a), but rather explores a range of tilt angles relative to the membrane’s normal (Figure 4b).

The overall gaussian distributions of the distances and angles measured here emphasize a continuous sampling of the distinct conformations, with no significant energy barriers hindering the transitions. Combined with the bi-directional tilting of each monomer (Supplementary Figure 1), the simulations indicate that the deformations occur transiently and with no preferred direction or conformation. Taken together, our results suggest that the experimentally resolved extended ACE2 structure is likely not a dominant conformation in solution, and the homodimer displays a large ensemble of conformations in the native state.

### ACE2-RBD interface remains stable despite the large ACE2 motions

Despite the dramatic flexibility of the ACE2 dimer, the RBD included in the RBD-bound model retained a large fraction of the native contacts with ACE2 throughout the simulations, with an average fraction of 0.87 ± 0.11 contacts. While the interface is thus overall stable, and the relatively small RBDs accompany the range of motion of ACE2, dividing the RBD-ACE2 interface into three interacting regions (comprising of the two RBD loop regions at the opposite sides of the dimer interface and the central region containing the two short b sheet strands, Figure 5a) we observe that the central region 2 contains the most stable contacts, while regions 1 and 3 at the extremities of the interface are less tightly bound and sample states with a smaller number of native contacts (Figure 5b). This rocking motion is in agreement with dynamics at the PD-RBD interface observed in other simulations (41, 42).

**Figure 5.**
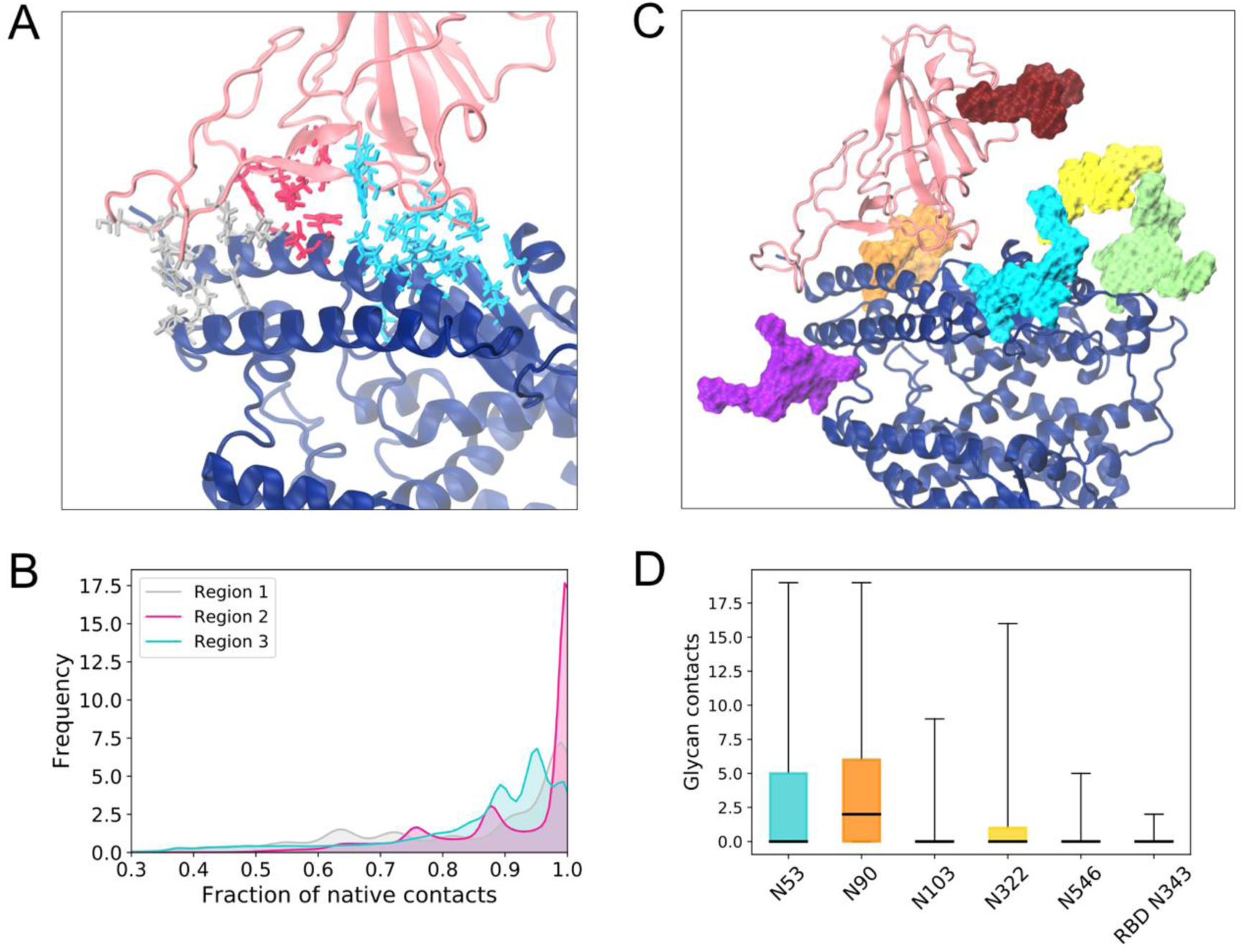
ACE2-RBD interactions. **(a)** Protein residue fraction of native contacts identified in the reference cryo-EM structure colored according to regions along the heterodimer interface (silver, magenta and cyan) and shown with licorice representation. ACE2 monomer shown with dark blue cartoons and RBD with pink cartoons. Glycans have been omitted from this panel for clarity. **(b)** Distribution of the fraction of native contacts in each of the interaction regions. Colors same as in (a). **(c)** Glycans in the ACE2-RBD interface, shown with surface representation with the following color scheme: N53 (cyan), N90 (orange), N103 (purple), N322 (yellow), N546 (lime) and N343 (dark red). **(d)** Box plot of number of glycan-protein contacts for the interface glycans shown in (c), using the same color scheme. Horizontal black lines indicate mean value, boxes extend to the lower and upper quartiles, and whiskers show the total range of the data.

In addition to protein interactions, five glycans in ACE2 are in close proximity to the RBD and have been suggested to play a role in S binding. In agreement with other studies (46, 63), N90 and to a lesser extent N322 of ACE2 establish contacts with RBD. Besides these glycans, we also find that N53 can form a large number of contacts with the RBD residues, while the RBD glycan N343 makes very few contacts with ACE2’s head protein residues (Figure 5c) or glycans (Supplementary Figure 3).

### N53 is involved in both homodimer and heterodimer contacts

Besides the ACE2-RBD heterodimer interface, we considered the interactions within ACE2 that could contribute to maintaining such a stable homodimer head interface despite the pronounced flexibility of the protein (Figures 2e and 3e). Experimental structures and simulations suggest that the majority of the protein contacts in the homodimer are located in the neck domain, with only two other interactions, in the form of hydrogen bonds, observed in the larger peptidase domain (16, 63). In agreement with these observations, we find that the dimer interface is mainly held together in the simulations by residues at the neck (Figure 6a for RBD-bound simulations and Supplementary Figure 4a for apo). Computation of the glycan-protein and glycan-glycan contacts enrich the characterization of the inter-and intra-monomer interactions and suggest that, while the eight ACE2 glycans form several contacts with protein residues located within the same monomer (Figure 6b), protein-glycan interactions with the opposite monomer are limited to N53 and N690 (Figure 6c). Additionally, N53 is the only glycan that can be found to form inter-monomer glycan-glycan contacts, established between the equivalent N53 copies in each of the monomers (Figure 6d and Supplementary Figure 5). Similarly to the lack of RBD effect on the ACE2 dimer flexibility, we find that the homodimer contact distributions are comparable between apo and RBD-bound states of ACE2 (Supplementary Figure 3).

**Figure 6.**
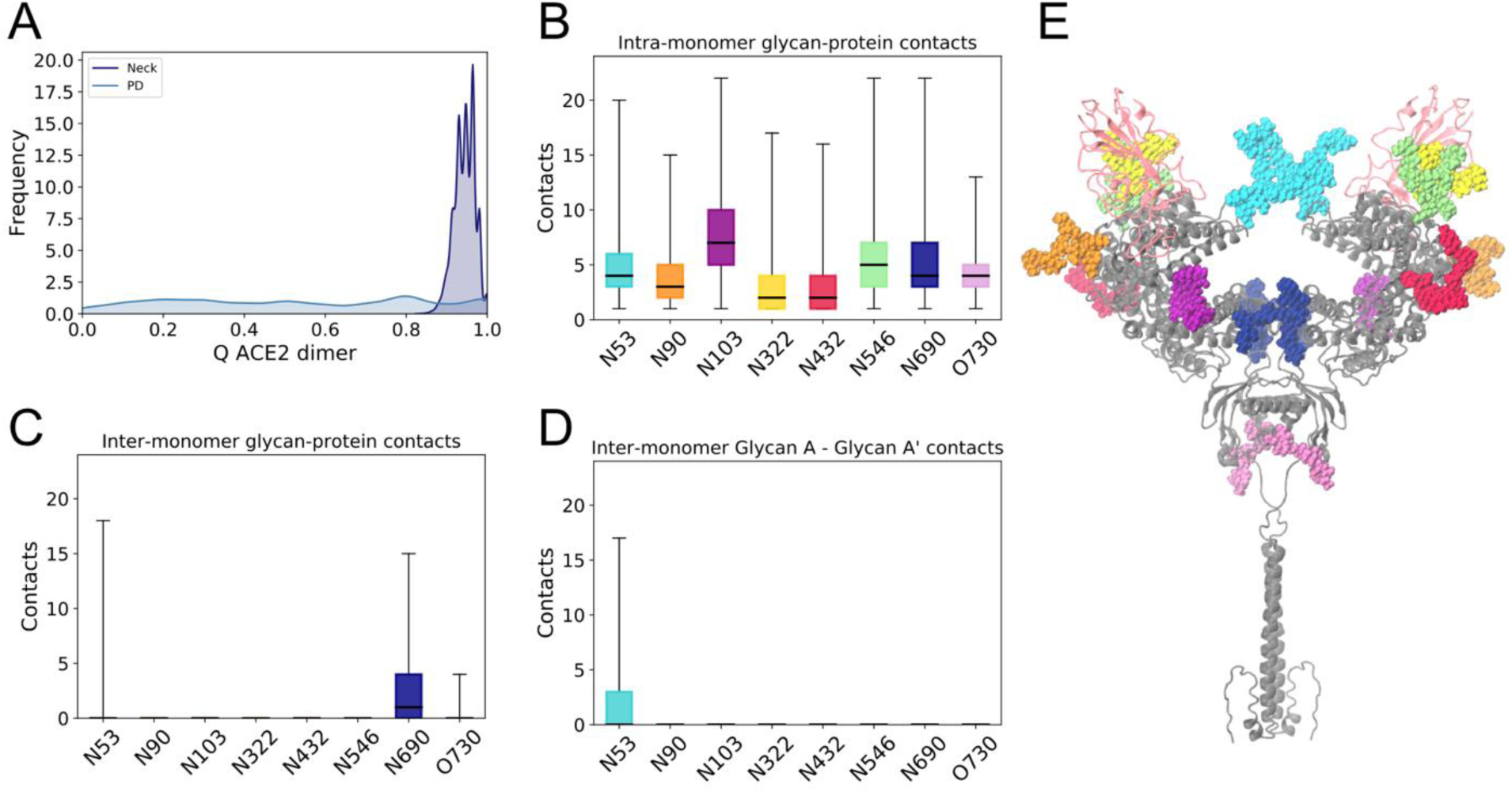
ACE2 homodimer contacts for RBD-bound simulations. **(a)** Fraction of native contacts between ACE2 monomers, considering only protein components of the glycoprotein. Neck and peptidase domain (PD) interacting regions are depicted separately. **(b)** Total glycan-protein interactions formed within each ACE2 monomer, per glycan. Horizontal black lines indicate mean value, boxes extend to the lower and upper quartiles, and whiskers show the total range of the data. **(c)** Glycan-protein contacts between glycans in one of the monomer and protein residues in the opposite monomer. **(d)** Glycan-glycan contacts between glycan in one of the monomers (glycan A) and its copy in the other monomer (glycan A’). **(e)** ACE2 dimer with glycans in van der Waals representation colored according to figures b-d. ACE2 protein dimer colored grey and RBDs in light pink.

Our systematic analysis of all glycan interactions thus indicate that N53 is the only glycan involved in both heterodimer (ACE2-RBD, Figure 5c) and homodimer (intra ACE2 dimer, Figure 6d) interactions. The opposite positions of the homodimer and RBD interfaces relative to N53 (Figure 5e) could suggest a competition for N53 contacts within these dimer interfaces. Indeed, we find that when a large number of N53-RBD contacts are formed, the N53 dimer interface is abrogated in the RBD-bound simulations, and vice versa (Supplementary Figure 6b). However, these interactions are not completely mutually exclusive as N53’ (the N53 glycan in monomer B) can be found involved in both inter-monomer and monomer-RBD interactions in Replica 1. This glycan’s flexibility probably plays a role in conferring a transient nature to the interactions, as the N53 homodimer contacts are not consistently formed even in the absence of the competing heterodimer in the apo ACE2 simulations (Supplementary Figure 6a). Thus, N53 can still be found highly solvent exposed in the apo state, suggesting optimal conformations for contact with an RBD partner and a role in S binding and infection.

Finally, as the large tilt motion of the ACE2 homodimer can bring the head domains in close contact to the membrane (Figure 1e), we also investigated the possible involvement of the glycans in interactions with the membrane polar heads. Although N103, N432 and N690 can establish up to 10 contacts at once with the membrane, we find that only O730, the glycan located closest to the membrane even in the elongated conformation, makes significant contacts with the lipids (Supplementary Figure 7), thus suggesting that ACE2’s glycans do not promote or stabilize the bent conformations.

## Discussion

All-atom simulations of apo and RBD-bound, full-length, membrane embedded ACE2 show a striking degree of fluctuation of the homodimer protein, which can be attributed to hinge motions of the large head domain relative to the transmembrane helices, and tilt of the transmembrane helices relative to the membrane’s normal. The head relative motion is due to the flexible linker region connecting the TM helix and the neck domain, while the TM helix motion points to a loose anchoring of ACE2 to the membrane. While the two (head and transmembrane) domains are internally stable, the flexibility of the connecting loop virtually results in a decoupling of these domains’ dynamics (Figure 7a), leading to sampling of conformations strikingly different than the experimentally-observed elongated structure (16). A high deformation propensity was also observed for the TM-neck linker upon normal model analysis of full-length ACE2 (71). These distinct conformations were likely only observed in the simulations due to the removal of B^0^AT1 from the co-complexed structure, since they seem to bind on opposite sides of the homodimer transmembrane interface and interact with the flexible linker (16). However, the expression profiles of ACE2 and B^0^AT1 suggests the likelihood of ACE2 existing in the apo state, especially in lung and heart tissues where B^0^AT1 is not expressed (48).

**Figure 7.**
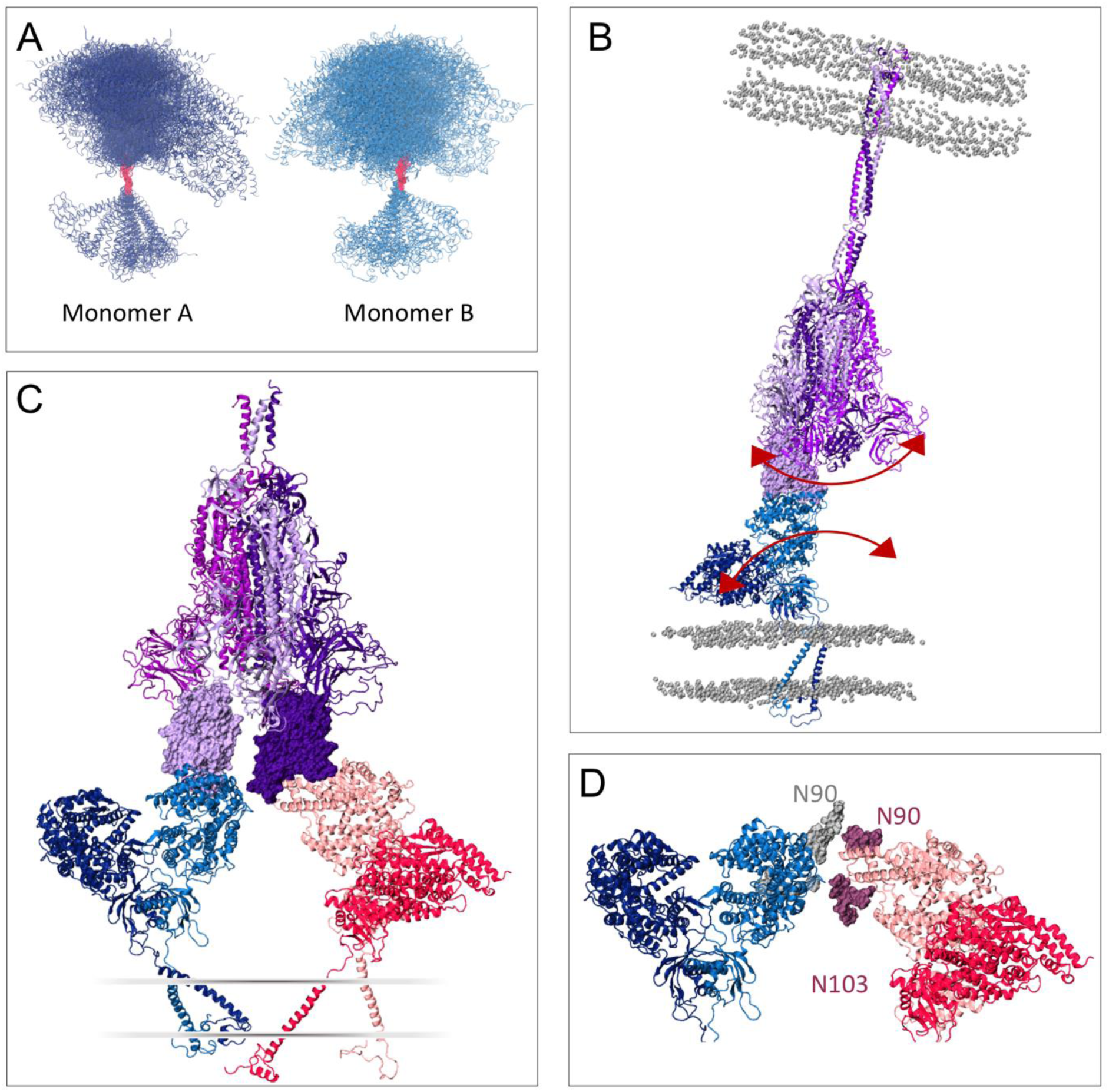
ACE2 flexibility’s impact on S interaction. **(a)** ACE2 monomer conformations taken from equally-spaced frames from the simulations, aligned via the flexible linker. **(b)** Proposed effect of ACE2’s flexibility on the spike’s dynamics, communicated through the ACE2-RBD complex. The three chains in the spike model are colored in different shades of purple, with the “up” RBD shown in light purple in surface representation. Phosphorus atoms from membrane’s lipid heads shown in grey in van der Waals representation. **(c)** Proposed complex of two ACE2 dimers bound to a single spike with two RBDs in the “up” conformation. ACE2 dimers shown in dark and light blue, and dark and light pink, respectively. RBDs shown in surface representation. A schematic of the membrane is indicated. (d) Detailed view of the ACE2 heads in (c), with N90 and N103 glycans highlighted.

Remarkably, the conformations of the ACE2 peptidase and neck domains remain stable throughout the simulations, and the homodimer heads retain their relative orientation despite the dramatic global homodimer motions. In a similar fashion, the RBDs included in the holo simulations remained tightly bound to ACE2 throughout the simulations, evidencing the high affinity between them. Glycan-glycan and glycan-protein interactions suggest that the ACE2 homodimer interface is maintained not only via protein interactions at the neck domain, but also inter-monomer contacts involving N53, at the top of the PD, and N690, closer to the neck. Interestingly, N53 also makes extensive contacts with RBD, suggesting a dual and possibly competing role between homodimer (intra-ACE2) and heterodimer (ACE2-RBD) interactions. This dual nature may be dependent on the length of the N53 glycan, but the glycosylation heterogeneity in ACE2 in general and in this position in particular (47) supports the likelihood of inter-monomer glycan interactions. Even in the absence of RBD, N53 alternates between being sequestered in homodimer contact and being extended and highly solvent accessible, suggesting a role in RBD binding to this glycan.

Due to the stability of the head domain and RBD interface in spite of ACE2 body motion, this large flexibility would remain invisible in studies that do not take the full length character of ACE2 into account, looking for instance at only the PD-RBD interactions. However, a recent cryo-EM study of S-ACE2 PD complex resolved a continuous swing motion of the ACE2 head-RBD relative to the S trimer body (44). These structural characterizations complement our analysis and suggest how the ACE2 motion would translate in the context of full-length spike. Additionally, it has been found from in-situ Cryo-EM and molecular dynamics simulations that the spike glycoprotein also exhibits conformational plasticity, with hinge motions at three different regions of the stalk trimer (43). Large dynamical variations thus seems to be a feature of these extracellular glycoproteins. The RBD rocking motion and S conformational variability have been proposed as mechanisms for immune evasion and efficient receptor search in the host cell (43, 44) but the similar rocking motion of ACE2 we observed also suggests a mechanical aspect to ACE2-S interaction. The process of S conformational transition upon binding to the receptor and cell fusion remains elusive, but ACE2’s intrinsic flexibility could promote a large swinging motion of the ACE2-S1 complex, providing a mechanical force for the approximation of the two membranes and shedding of S1 towards fusion of the S2 domains into the receptor cell (Figure 7b).

Finally, one can speculate that the flexibility of the host receptor might allow the accommodation of more than one ACE2 dimer bound to a single S with two or more RBDs in the up conformation. A high efficiency of ACE2 usage was suggested to contribute to SARS-CoV transmissibility(15, 72), and thus could be at play for SARS-CoV-2 as well. To investigate this possibility, we extracted a range of ACE2 conformations from the RBD-bound simulations covering different tilt angles, and explored the alignment of these structures to a “three RBD-up” spike model. Indeed, we find that two ACE2 dimers can sterically be accommodated by a single spike, with inter-dimer backbone distances no smaller than 10 Å (Figure 7d). The flexibility of the homodimers could potentially allow for even three ACE2’s per S, opening the possibility of multi-receptor usage by the spike glycoprotein for host cell infection. Explicitly considering the glycans in this aligned model evidences that N103 and especially N90 are in close proximity to the neighboring ACE2 dimer (Figure 7c). Interestingly, it is known that disruption of the N90 glycosylation motif due to mutations leads to increased S-ACE2 binding affinity (72–74), and these observations can thus provide the structural basis for the negative effect of N90 on RBD binding.

## Conclusions

All-atom molecular dynamics simulations of the full-length ACE2 inserted in a mammalian-inspired lipid membrane uncover a significant degree of flexibility of the ACE2 homodimer with consequences for S-ACE2 interaction and SARS-CoV-2 infection, and suggest the structural basis for glycan N90’s negative effect on RBD binding. Additionally, we identify the involvement of glycan N53 in ACE2 homodimer and ACE2-RBD heterodimer contacts. Taken together, our findings shed further light onto the mechanisms of viral binding and cell entry required for rational design of effective SARS-CoV-2 therapeutics.

## Author Contributions

E.P.B., L.C. and Z.G. built model of the ACE2 and RBD. A.C.D. built the membrane bilayer. E.P.B. performed the MD simulations and analysis. Y.Z. built the 3-up RBD, L.F. converted the spike and glycans from Charmm to AMBER and GLYCAM force fields, L.F. performed the MD of the spike structures, K.F. did the structure grafting. R.E.A. and C.S. designed and oversaw the research project. E.P.B. wrote the paper with contributions from all authors.

## Acknowledgments

The authors declare no conflict of interest. This work was supported by NIH GM132826, NSF RAPID MCB-2032054, an award from the RCSA Research Corp., and a UC San Diego Moore’s Cancer Center 2020 SARS-COV-2 seed grant. LC is funded by a Visible Molecular Cell Consortium fellowship.

